# Catenin signaling controls phrenic motor neuron development and function during a narrow temporal window

**DOI:** 10.1101/2023.01.18.524559

**Authors:** Alicia N. Vagnozzi, Matthew T. Moore, Raquel López de Boer, Aambar Agarwal, Niccolò Zampieri, Lynn T. Landmesser, Polyxeni Philippidou

## Abstract

Phrenic Motor Column (PMC) neurons are a specialized subset of motor neurons (MNs) that provide the only motor innervation to the diaphragm muscle and are therefore essential for survival. Despite their critical role, the mechanisms that control phrenic MN development and function are not well understood. Here, we show that catenin-mediated cadherin adhesive function is required for multiple aspects of phrenic MN development. Deletion of *β*- and *γ*-catenin from MN progenitors results in perinatal lethality and a severe reduction in phrenic MN bursting activity. In the absence of catenin signaling, phrenic MN topography is eroded, MN clustering is lost and phrenic axons and dendrites fail to grow appropriately. Despite the essential requirement for catenins in early phrenic MN development, they appear to be dispensable for phrenic MN maintenance, as catenin deletion from postmitotic MNs does not impact phrenic MN topography or function. Our data reveal a fundamental role for catenins in PMC development and suggest that distinct mechanisms are likely to control PMC maintenance.

## Introduction

Breathing is an essential motor behavior that is required for survival. In mammals, contraction of the diaphragm muscle is critical for bringing oxygenated air into the lungs during inspiration (Greer, 2012). Diaphragm contractions are mediated by a specialized subset of motor neurons (MNs), Phrenic Motor Column (PMC) neurons that reside in the cervical spinal cord and project their axons along the phrenic nerve in the thoracic cavity to reach the diaphragm. Phrenic MNs exhibit distinct properties from other MN subtypes, including tight clustering and stereotyped axonal and dendritic morphologies. While they are derived from a common MN progenitor domain, phrenic MNs acquire their unique features through the activity of a selective transcriptional program that distinguishes them from other MNs (Chaimowicz et al., 2019; Machado et al., 2014; Philippidou et al., 2012; Vagnozzi et al., 2020). Phrenic-specific transcription factors (TFs) initiate and maintain the expression of a distinct set of genes, including a unique combination of cell surface adhesion molecules (Machado et al., 2014; Vagnozzi et al., 2020). While many of these molecular markers show specific and sustained expression in phrenic MNs, their functions in phrenic MN development and maintenance have not been tested.

We previously identified a distinct combinatorial cadherin code that defines phrenic MNs, which includes both the broadly expressed type I N-cadherin and a subset of specific type II cadherins (Vagnozzi et al., 2020). Cadherins are calcium-dependent transmembrane cell adhesion molecules that interact with cytosolic catenin proteins to induce changes in the actin cytoskeleton, thus regulating many neuronal processes such as migration, topography and morphology (Seong et al., 2015). For example, cadherins regulate cortical neuron migration (Martinez-Garay, 2020), hippocampal dendritic growth and branching (Bekirov et al., 2008; Esch et al., 2000; Yu and Malenka, 2003), as well as dendrite morphogenesis and arborization within the visual and olfactory systems (Duan et al., 2018; Hirano and Takeichi, 2012; Masai et al., 2003; Riehl et al., 1996; Tanabe et al., 2006; Zhu and Luo, 2004). In the spinal cord, cadherins engage β- and γ-catenins to establish the segregation and settling position of MN cell bodies (Demireva et al., 2011; Dewitz et al., 2019; Dewitz et al., 2018; Price et al., 2002). β-catenin is required in muscle for neuromuscular junction (NMJ) formation and function, however it appears to act redundantly with γ-catenin in MNs, as only joint β- and γ-catenin inactivation leads to disorganization of MN subtypes, including PMC neurons (Demireva et al., 2011; Li et al., 2008; Vagnozzi et al., 2020). However, it is unknown whether β- and γ-catenins have additional roles in phrenic MN development and function, and whether they continue to be required after initial PMC topography has been established.

Here, we show that catenin activity is required for proper respiratory behavior and robust respiratory output. After MN-specific deletion of *β*- and *γ*-catenin, mice display severe respiratory insufficiency, gasp for breath, and die within hours of birth. Using phrenic nerve recordings, we determined that catenins are crucial for respiratory motor output, as MN-specific catenin inactivation leads to a striking decrease in phrenic MN activity. We further show that catenins are required to establish phrenic MN cell body settling position, as well as PMC axonal and dendritic morphology. Finally, we show that catenins are only required for PMC development and function during a narrow developmental window, as catenin deletion from postmitotic MNs does not impact respiratory output. Our data demonstrate a fundamental role for the cadherin-catenin cell adhesion complex in phrenic MN development and respiratory function and indicate that distinct pathways likely act to maintain PMC function.

## Materials and Methods

### Mouse genetics

The *β*-catenin (Brault et al., 2001) and *γ*-catenin (Demireva et al., 2011) alleles, *Olig2::Cre* (Dessaud et al., 2007) and *ChAT::Cre* (Lowell et al., 2006) lines were generated as previously described and maintained on a mixed background. Mouse colony maintenance and handling was performed in compliance with protocols approved by the Institutional Animal Care Use Committee of Case Western Reserve University. Mice were housed in a 12-hour light/dark cycle in cages containing no more than five animals at a time.

### Immunohistochemistry and in situ hybridization

Immunohistochemistry was performed as previously described (Philippidou et al., 2012; Vagnozzi et al., 2020), on tissue fixed for 2 hours in 4% paraformaldehyde (PFA) and cryosectioned at 16μm. Wholemounts of diaphragm muscles were stained as described (Philippidou et al., 2012). The following antibodies were used: goat anti-Scip (1:5000; Santa Cruz Biotechnology, RRID:AB_2268536), mouse anti-Islet1/2 (1:1000, DSHB, RRID:AB_2314683) (Tsuchida et al., 1994), rabbit anti-neurofilament (1:1000; Synaptic Systems, RRID:AB_887743), rabbit anti-synaptophysin (1:250, Thermo Fisher, RRID:AB_10983675), and α-bungarotoxin Alexa Fluor 555 conjugate (1:1000; Invitrogen, RRID:AB_2617152). Images were obtained with a Zeiss LSM 800 confocal microscope and analyzed with Zen Blue, ImageJ (Fiji), and Imaris (Bitplane). Phrenic MN number was quantified using the Imaris “spots” function to detect cell bodies that coexpressed high levels of Scip and Isl1/2 in a region of interest limited to the left and right sides of the ventral spinal cord.

### DiI tracing

For labeling of phrenic MNs, crystals of carbocyanine dye, DiI (Invitrogen, #D3911) were pressed onto the phrenic nerves of eviscerated embryos at e18.5, and the embryos were incubated in 4% PFA at 37°C in the dark for 4-5 weeks. Spinal cords were then dissected, embedded in 4% low melting point agarose (Invitrogen) and sectioned using a Leica VT1000S vibratome at 100 to 150μm.

### Positional analysis

MN positional analysis was performed as previously described (Dewitz et al., 2019; Dewitz et al., 2018). MN positions were acquired using the “spots” function of the imaging software Imaris (Bitplane) to assign x and y coordinates. Coordinates were expressed relative to the midpoint of the spinal cord midline, defined as position x=0, y=0. To account for experimental variations in spinal cord size, orientation, and shape, sections were normalized to a standardized spinal cord whose dimensions were empirically calculated at e13.5 (midline to the lateral edge=390μm). We analyzed every other section containing the entire PMC (20-30 sections in total per embryo).

### Dendritic orientation analysis

For the analysis of dendritic orientation, we superimposed a radial grid divided into eighths (45 degrees per octant) centered over phrenic MN cell bodies spanning the entire length of the dendrites. We drew a circle around the cell bodies and deleted the fluorescence associated with them. Fiji (ImageJ) was used to calculate the fluorescent intensity (IntDen) in each octant which was divided by the sum of the total fluorescent intensity to calculate the percentage of dendritic intensity in each area.

### Electrophysiology

Electrophysiology was performed as previously described (Vagnozzi et al., 2020). Mice were cryoanesthetized and rapid dissection was carried out in 22-26°C oxygenated Ringer’s solution. The solution was composed of 128mM NaCl, 4mM KCl, 21mM NaHCO_3_, 0.5mM NaH_2_PO_4_, 2mM CaCl_2_, 1mM MgCl_2_, and 30mM D-glucose and was equilibrated by bubbling in 95% O_2_/5% CO_2_. The hindbrain and spinal cord were exposed by ventral laminectomy, and phrenic nerves exposed and dissected free of connective tissue. A transection at the pontomedullary boundary rostral to the anterior inferior cerebellar artery was used to initiate fictive inspiration. Electrophysiology was performed under continuous perfusion of oxygenated Ringer’s solution from rostral to caudal. Suction electrodes were attached to phrenic nerves just proximal to their arrival at the diaphragm. The signal was band-pass filtered from 10Hz to 3kHz using AM-Systems amplifiers (Model 3000), amplified 5,000-fold, and sampled at a rate of 50kHz with a Digidata 1440A (Molecular Devices). Data were recorded using AxoScope software (Molecular Devices) and analyzed in Spike2 (Cambridge Electronic Design). Burst duration and burst activity were computed from 4-5 bursts per mouse, while burst frequency was determined from 10 or more minutes of recording time per mouse. Burst activity was computed by rectifying and integrating the traces with an integration time equal to 2 seconds, long enough to encompass the entire burst. The maximum amplitude of the rectified and integrated signal was then measured and reported as the total burst activity.

### Plethysmography

Conscious, unrestrained P0 mice were placed in a whole body, flow-through plethysmograph (emka) attached to a differential pressure transducer (emka). We modified 10ml syringes to use as chambers, as smaller chambers increase signal detection in younger mice. Experiments were done in room air (79% nitrogen, 21% oxygen). Mice were placed in the chamber for 30 seconds at a time, for a total of three to five times, and breathing parameters were recorded. Mice were directly observed to identify resting breaths. At least ten resting breaths were analyzed from every mouse. Data are presented as fold control, where the control is the average of 2 littermates in normal air.

### Experimental design and statistical analysis

For all experiments a minimum of three embryos per genotype, both male and female, were used for all reported results unless otherwise stated. The Shapiro-Wilk test was used to determine normality. All data, with the exception of electrophysiology burst activity data in figures 2 and 6, showed normal distribution and p-values were calculated using unpaired, two-tailed Student’s *t* test. Burst activity data in figures 2 and 6 showed non-normal distribution and p-values were calculated using the Mann-Whitney test. p < 0.05 was considered to be statistically significant, where * p< 0.05, ** p < 0.01, *** p < 0.001, and **** p < 0.0001. Data are presented as box and whisker plots with each dot representing data from one mouse unless otherwise stated. Small open squares in box and whisker plots represent the mean.

**Figure 1.**
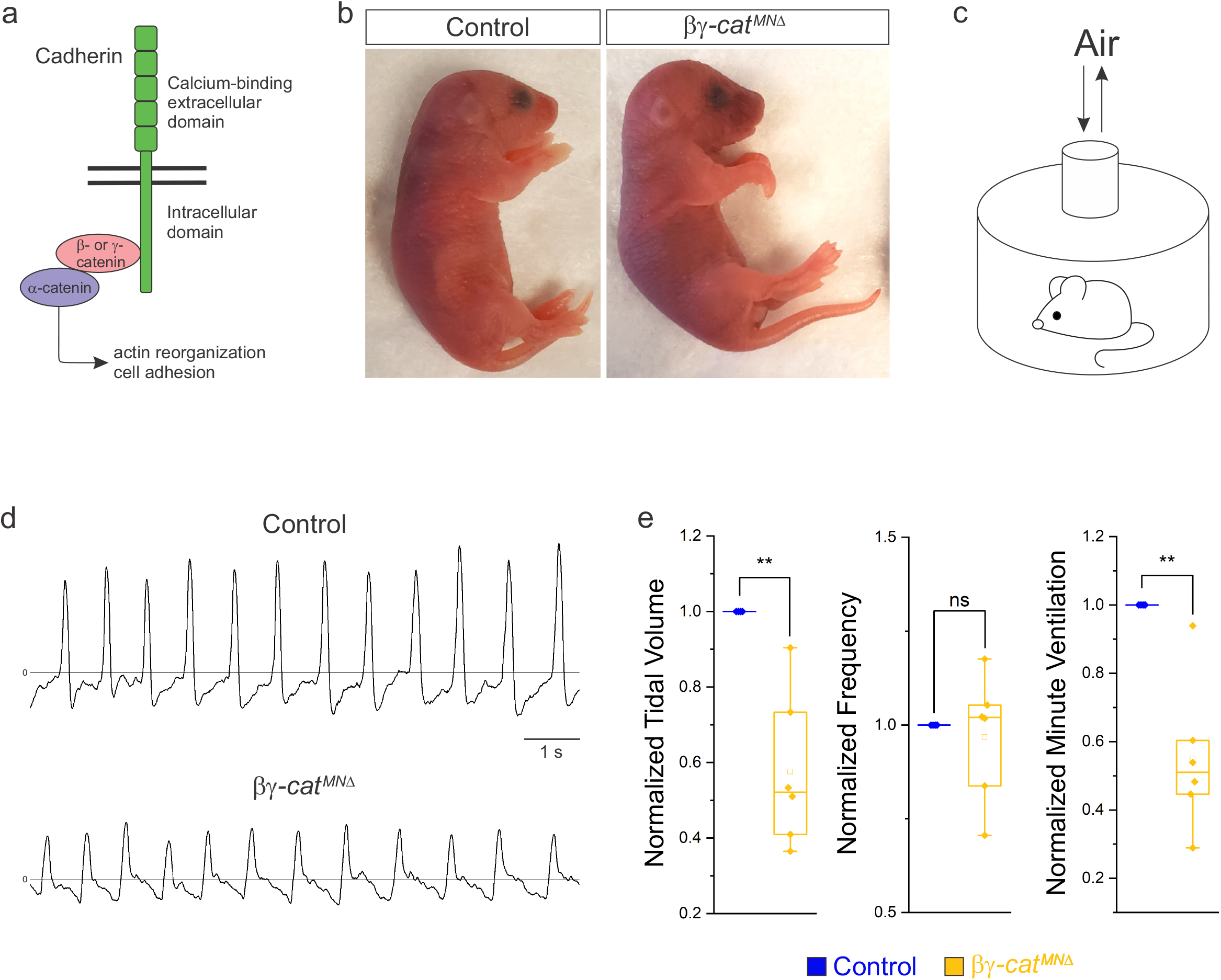
Catenin signaling is required for survival and proper respiratory behavior. **a)** B- and γ-catenin are obligate intracellular factors required for cadherin-mediated cell adhesive function. We utilized inactivation of β- and γ-catenin in MNs (*β-catenin flox/flox*; *γ-catenin flox/flox*; *Olig2::Cre*, referred to as *βγ*-*cat*^*MN*Δ^ mice) as a strategy to define the function of cadherin signaling in phrenic MNs. **b)** Appearance of P0 control and *βγ*-*cat*^*MN*Δ^ mice. *Bγ*-*cat*^*MN*Δ^ mice are cyanotic and die shortly after birth. **c)** Experimental setup for whole body plethysmography experiments. **d)** Representative 10 second traces in room air from control and *βγ*-*cat*^*MN*Δ^ mice at P0. **e)** *βγ*-*cat*^*MN*Δ^ mice display reduced tidal volume resulting in a 45% reduction in overall ventilation (n=4 control, n=6 *βγ*-*cat*^*MN*Δ^ mice).

**Figure 2.**
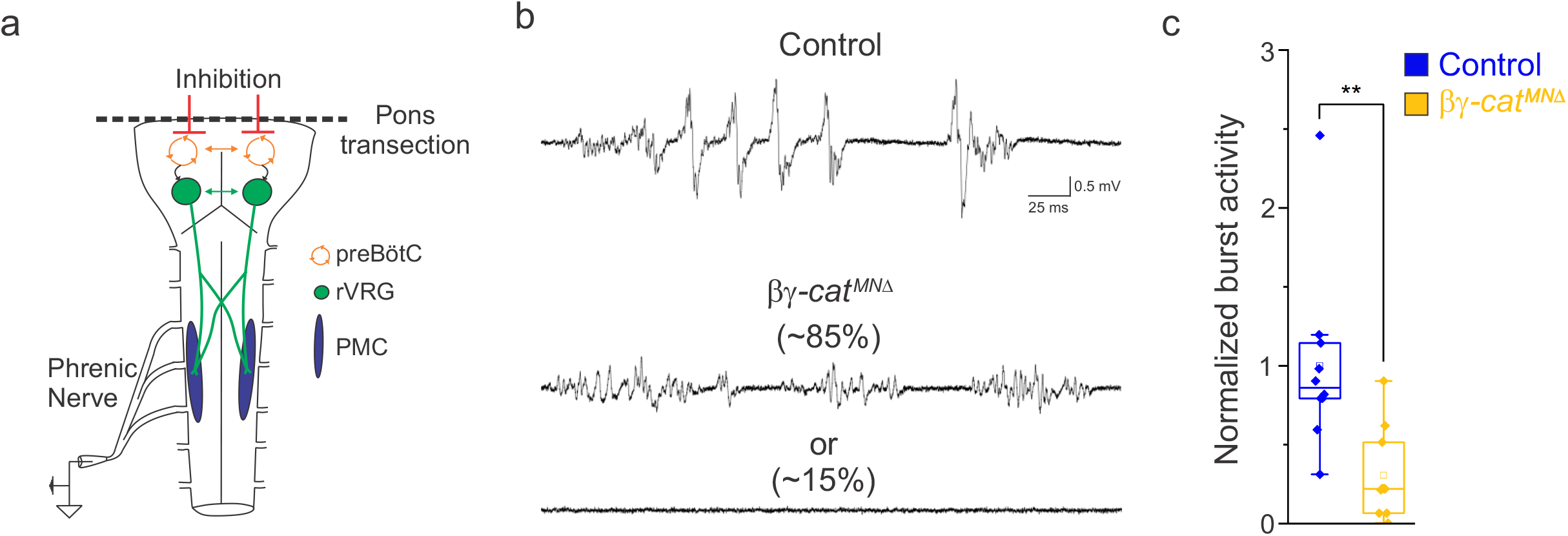
Catenin signaling controls phrenic MN activation. **a)** Schematic of brainstem-spinal cord preparation, which displays fictive inspiration after removal of the pons. Suction electrode recordings were taken from the phrenic nerve in the thoracic cavity at e18.5/P0. **b)** Enlargement of single respiratory bursts reveals a reduction in burst amplitude and overall activity in *βγ*-*cat*^*MN*Δ^ mice. Partial (initial 350ms) bursts are shown. While 85% of *βγ*-*cat*^*MN*Δ^ mice display respiratory bursts, 15% show no bursts throughout the recording period. **c)** *βγ*-*cat*^*MN*Δ^ mice exhibit nearly 70% reduction in burst activity (n=10 control, n=10 *βγ*-*cat*^*MN*Δ^ mice).

## Results

### Catenin signaling is required for survival, proper respiratory behavior and phrenic MN activation

Phrenic MNs express a distinct combinatorial cadherin code (Machado et al., 2014; Vagnozzi et al., 2020), but the collective contribution of these molecules to phrenic MN development, maintenance and function has not been established. We previously found that phrenic MNs express the type I *N-cadherin* (*N-cad*) and the type II cadherins *Cdh6, 9, 10, 11*, and *22* (Vagnozzi et al., 2020). To investigate the role of cadherin signaling in phrenic MN development, we eliminated cadherin signaling in MN progenitors by inactivating *β*- and *γ*-catenin using a *Olig2::Cre* promoter (*β-catenin flox/flox; γ-catenin flox/flox; Olig2::Cre*, referred to as *βγ*-*cat*^*MN*Δ^ mice). β- and γ-catenin are obligate intracellular factors required for cadherin-mediated cell adhesive function and catenin deletion enables us to interrogate the full repertoire of cadherin actions in phrenic MNs (Figure 1a). We find that *βγ*-*cat*^*MN*Δ^ mice are born alive but die within 24 hours of birth and often display severe flexion of the wrist joint (Figure 1b).

In order to assess breathing in *βγ*-*cat*^*MN*Δ^ mice, we utilized unrestrained whole body flow-through plethysmography at postnatal day (P)0 (Figure 1c). We found that *βγ*-*cat*^*MN*Δ^ mice have a 45% reduction in tidal volume (the amount of air inhaled during a normal breath), while respiratory frequency is not affected (Figure 1d-e). This results in an average 45% reduction in overall air drawn into the lungs per minute (minute ventilation, figure 1e), indicating *βγ*-*cat*^*MN*Δ^ mice likely die from respiratory failure. Our findings indicate that catenin signaling is necessary for proper respiratory behavior and survival. To further examine respiratory circuitry intrinsic to the brainstem and spinal cord, we performed suction recordings of the phrenic nerve in isolated brainstem-spinal cord preparations (Figure 2a). These preparations display fictive inspiration after the removal of inhibitory networks in the pons via transection, and thus represent a robust model to interrogate circuit level changes. We examined whether catenin deletion impacts circuit output at embryonic day (e)18.5/P0, shortly before *βγ*-*cat*^*MN*Δ^ mice die. We observed a striking reduction in the activation of phrenic MNs in *βγ*-*cat*^*MN*Δ^ mice (Figure 2b). While bursts in control mice exhibit large peak amplitude, bursts in *βγ*-*cat*^*MN*Δ^ mice were either of very low amplitude (~85%) or non-detectable (~15%). After rectifying and integrating the traces, we found a nearly 70% decrease in total burst activity in *βγ*-*cat*^*MN*Δ^ mice (Figure 2c). Our data indicate that cadherin signaling is imperative for robust activation of phrenic MNs during inspiration.

### Catenins establish phrenic MN topography and organization

What accounts for the loss of phrenic MN activity in *βγ*-*cat*^*MN*Δ^ mice? We asked whether early phrenic MN specification, migration and survival are impacted after catenin inactivation. We acquired transverse spinal cord sections through the entire PMC at e13.5 and stained for the phrenic-specific TF Scip and the MN-specific TF Isl1/2, to label all phrenic MNs. We found a cluster of Scip+ MNs in the ventral cervical spinal cord of both control and *βγ*-*cat*^*MN*Δ^ mice, indicating that early phrenic MN specification is unperturbed (Figure 3a). However, we observed a significant reduction in phrenic MN numbers in *βγ*-*cat*^*MN*Δ^ mice, indicating that catenin signaling is necessary for phrenic MN survival (Figure 3b).

**Figure 3.**
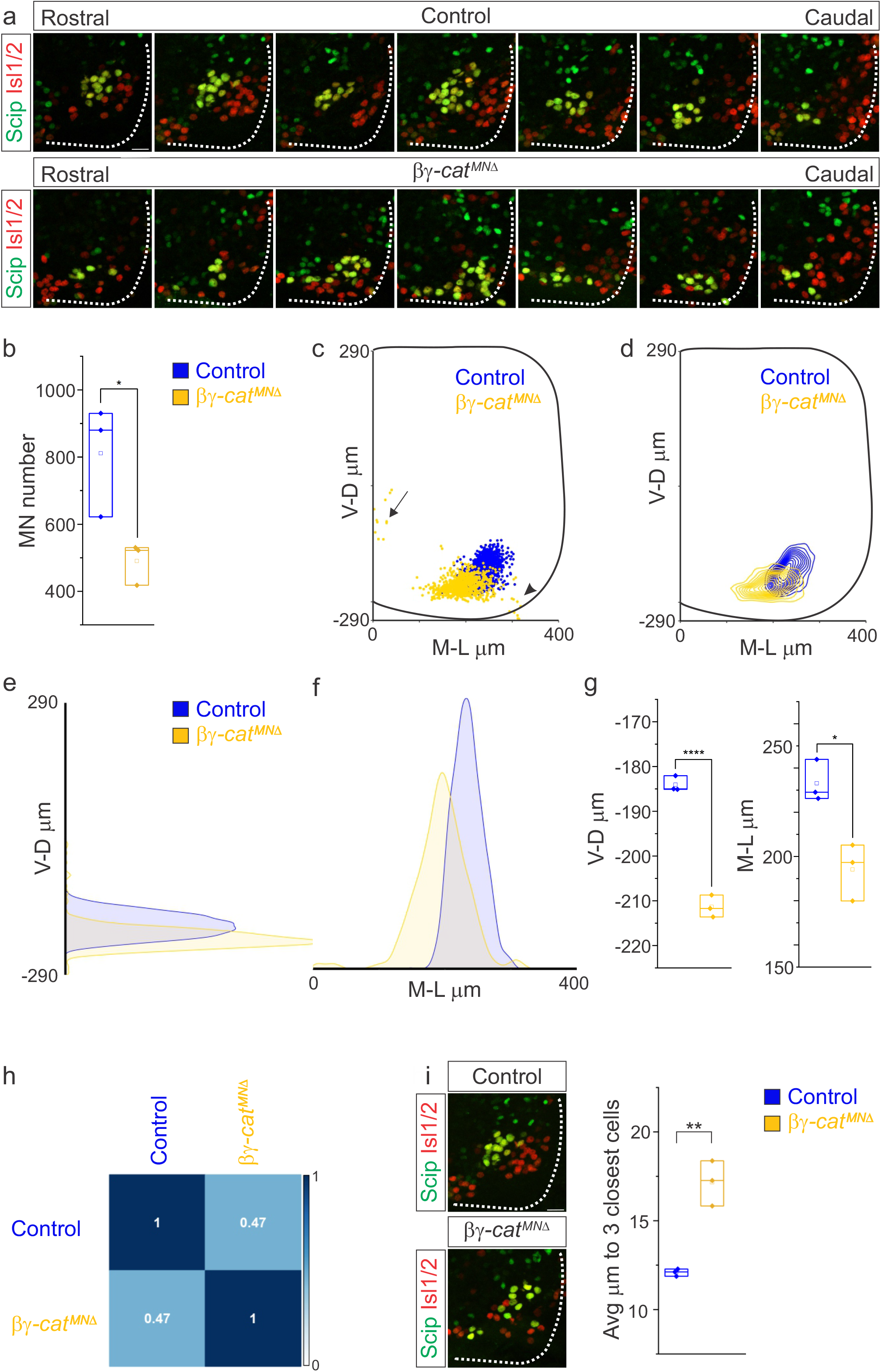
Catenins establish phrenic MN topography and organization. **a)** Rostral to caudal distribution of e13.5 phrenic MN cell bodies (yellow, defined by the expression of Scip in green and Isl1/2 in red) in control and *βγ*-*cat*^*MN*Δ^ mice. While phrenic MN cell bodies in control mice gradually shift towards more ventral positions at caudal locations, phrenic cell bodies are located ventrally in *βγ*-*cat*^*MN*Δ^ mice even at rostral levels. In addition, cell bodies in *βγ*-*cat*^*MN*Δ^ mice appear less clustered. **b)** Reduction in phrenic MN number in *βγ*-*cat*^*MN*Δ^ mice. **c)** Reconstructed distribution of cell bodies in control and *βγ*-*cat*^*MN*Δ^ mice. Occasional cell bodies remain near the progenitor zone in *βγ*-*cat*^*MN*Δ^ mice (arrow), while others seem to be dragged out of the spinal cord by their axon (arrowhead). **d)** Contour density plot of phrenic cell body position in control and *βγ*-*cat*^*MN*Δ^ mice at e13.5. V-D μm; ventrodorsal position, M-L μm; mediolateral position. (0,0) represents the center of the spinal cord in both dimensions. **e-f)** Density plots of ventrodorsal (e) and mediolateral (f) cell body position in control and *βγ*-*cat*^*MN*Δ^ mice. **g)** Quantification of ventrodorsal and mediolateral position, showing significant shifts in *βγ*-*cat*^*MN*Δ^ mice. **h)** Correlation analysis of phrenic MN positional coordinates in control and *βγ*-*cat*^*MN*Δ^ mice. 0 is no correlation, while 1 is a perfect correlation. **i)** Phrenic MNs have an increased average distance to their neighboring phrenic MNs in *βγ*-*cat*^*MN*Δ^ mice, indicating loss of clustering. Scale bar= 25μm.

While phrenic MNs are normally distributed along the rostrocaudal axis, we found that they sometimes show migratory defects, where several phrenic MNs remain close to the midline instead of fully migrating (Figure 3c, arrow), and also appear to shift both ventrally and medially in *βγ*-*cat*^*MN*Δ^ mice. To quantitate PMC cell body position, each phrenic MN was assigned a cartesian coordinate, with the midpoint of the spinal cord midline defined as (0,0). *Bγ*-*cat*^*MN*Δ^ mice displayed a significant shift in phrenic MN cell body position, with cell bodies shifting ventrally towards the edge of the spinal cord and towards the midline (Figure 3c-f). We quantified the average ventrodorsal and mediolateral phrenic MN position per embryo and found a significant change in phrenic MN position in both axes in *βγ*-*cat*^*MN*Δ^ mice (Figure 3g). Correlation analysis indicated that control and *βγ*-*cat*^*MN*Δ^ mice are dissimilar from each other (r=0.47, Figure 3h), indicating that catenins establish phrenic MN coordinates during development.

In addition to changes in cell body position, we also noticed that phrenic MNs appear to lose their tight clustering in *βγ*-*cat*^*MN*Δ^ mice. PMC clustering is thought to be critical for the proper development of the respiratory system because it facilitates recruitment of motor units through electrical coupling in the embryo to compensate for weak inspiratory drive (Greer and Funk, 2005). In order to determine PMC clustering defects, we used Imaris to measure the average distance between phrenic MNs. We found that *βγ*-*cat*^*MN*Δ^ mice had a nearly 50% increase in the average distance between neighboring cells (Figure 3i), indicating that the cadherin/catenin adhesive complex contributes to the formation of a tightly clustered phrenic motor column.

### Catenins are required for phrenic MN dendrite and axon growth

Since cadherin/catenin signaling is imperative for phrenic MN organization, we also asked whether any other aspects of phrenic MN development, such as dendritic and axon growth, might rely on catenin actions. We examined dendritic orientation in control and *βγ*-*cat*^*MN*Δ^ mice by injecting the lipophilic dye diI into the phrenic nerve at e18.5 (Figure 4a). DiI diffuses along the phrenic nerve to label both PMC cell bodies and dendrites. Consistent with our earlier observations, we found that phrenic cell bodies are often scattered in *βγ*-*cat*^*MN*Δ^ mice. Interestingly, even phrenic MNs that are significantly displaced correctly project along the phrenic nerve (arrows, figure 4a), indicating that changes in cell body position do not impact axon trajectory choice. In control mice, phrenic MN dendrites branch out in dorsolateral to ventromedial directions; in *βγ*-*cat*^*MN*Δ^ mice, however, they exhibit stunted growth, defasciculation and a failure to extend in the dorsolateral direction (Figure 4a).

**Figure 4.**
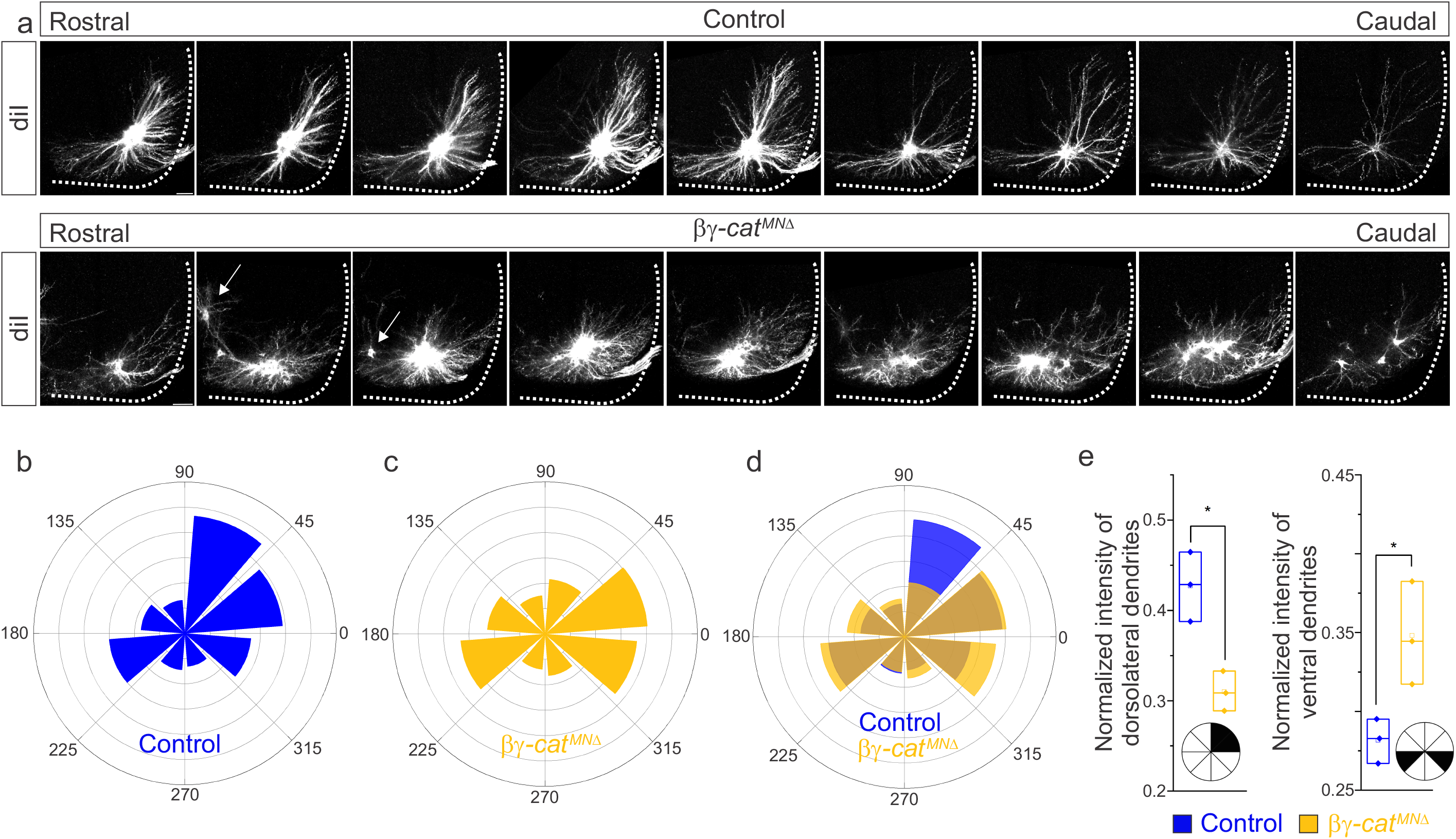
Catenins control phrenic MN dendritic growth and orientation. **a)** Rostral to caudal extent of phrenic MN dendrites, as revealed by diI injections into the phrenic nerve in control and *βγ*-*cat*^*MN*Δ^ mice. Scale bar= 100μm. **b-d)** Radial plot of the normalized fluorescent intensity in each octant in control (b, d) and *βγ*-*cat*^*MN*Δ^ (c-d) mice. Zero degrees represents a line through the center of the phrenic MN cell bodies that is perpendicular to the midline. Dendrites in *βγ*-*cat*^*MN*Δ^ mice shift ventrally, with a loss of dorsolateral projections. **e)** Quantification of the proportion of dendritic fluorescent intensity from 0 to 90 degrees (left, dorsolateral) and from 180 to 225 degrees and 315 to 360 degrees (right, ventral) in control and *βγ*-*cat*^*MN*Δ^ mice.

To quantify these changes, we superimposed a radial grid divided into octants onto the dendrites and measured the fluorescent intensity in each octant after removal of any fluorescence associated with the cell bodies. Zero degrees was defined by a line running perpendicular from the midline through the center of cell bodies. In control mice, the majority of dendrites project in the dorsolateral direction (0-90 degrees), representing 40-45% of the overall dendritic intensity (Figure 4b, d, e). Ventrally projecting dendrites (180-225 degrees and 315-360 degrees) were also prominent, giving rise to nearly 30% of the overall dendritic intensity (Figure 4b, d, e). We found that catenin deletion resulted in a striking reduction in dorsolateral dendrites, together with a significant increase in ventral dendrites (Figure 4c-e), indicating that cadherins are necessary for establishing phrenic MN dendritic orientation. These changes in phrenic dendritic topography in *βγ*-*cat*^*MN*Δ^ mice may impact their targeting by respiratory populations in the brainstem, leading to the reduction in phrenic MN activation observed.

We then asked whether catenins might also play an analogous role in phrenic axon extension and arborization. We examined diaphragm innervation in control and *βγ*-*cat*^*MN*Δ^ mice at e18.5. We found that *βγ*-*cat*^*MN*Δ^ mice display a lack of innervation in the ventral diaphragm (Figure 5a, arrow), while the parts of the diaphragm that are innervated show a reduction in terminal arborization complexity (Figure 5a, star). Quantitation of total phrenic projections revealed a significant reduction in overall diaphragm innervation (Figure 5b-c). Collectively our data point to a catenin requirement for phrenic MN topography and axon and dendrite arborization, suggesting that these changes in early phrenic MN development lead to loss of phrenic MN activity and perinatal lethality due to respiratory insufficiency.

**Figure 5.**
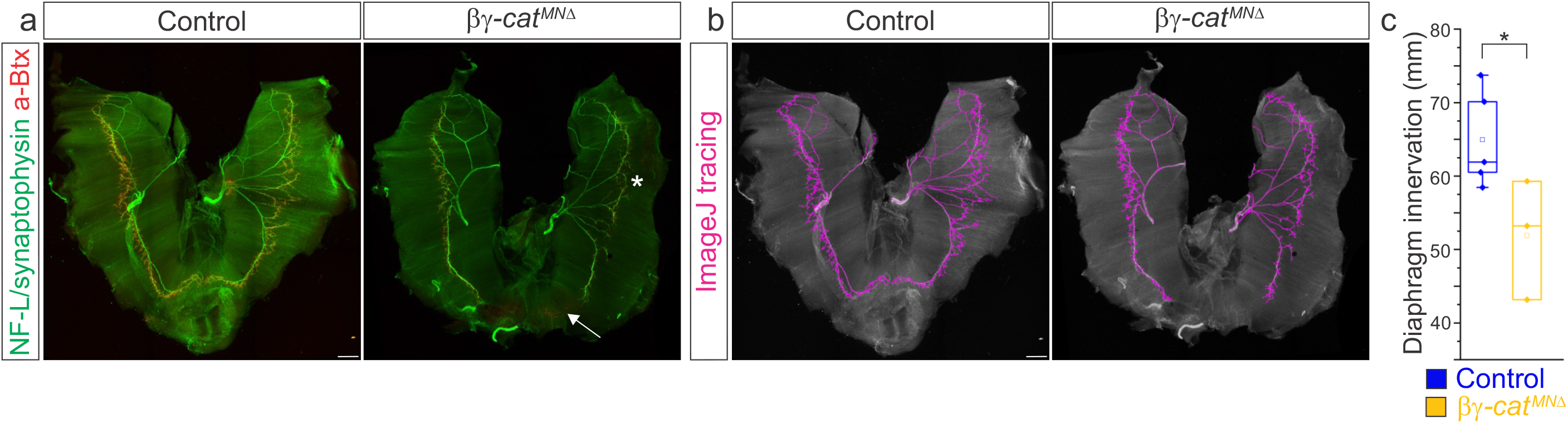
Catenins regulate phrenic MN axonal arborization. **a)** Diaphragm innervation in control and *βγ*-*cat*^*MN*Δ^ mice. *βγ*-*cat*^*MN*Δ^ mice display a reduction in ventral diaphragm innervation (arrow) and arborization complexity (star) at e18.5. Motor axons are labeled in green (combination of neurofilament light chain/synaptophysin) and neuromuscular junctions in red (α-bungarotoxin, btx). Scale bar= 500μm. **b)** Phrenic projections were traced and quantified in ImageJ. **c)** Quantification of diaphragm innervation in control and *βγ*-*cat*^*MN*Δ^ mice.

### A narrow temporal requirement for catenin signaling in phrenic MN topography and function

Given the essential role for catenins in early phrenic MN development, we wanted to further understand the temporal dynamics of cadherin signaling, and asked whether sustained catenin expression is required for maintenance of the respiratory circuit. We used a *ChAT::Cre* promoter to specifically delete *β*- and *γ-catenin* in postmitotic MNs (*β-catenin flox/flox*; *γ-catenin flox/flox*; *ChAT::Cre*, referred to as *βγ*-*cat*^*ChATMN*Δ^ mice). *βγ*-*cat*^*ChATMN*Δ^ mice survive to adulthood and do not display respiratory insufficiency or gasping at birth. We first assessed cell body position and found no changes between control and *βγ*-*cat*^*ChATMN*Δ^ mice (Figure 6a). Injecting diI into the phrenic nerve also revealed no differences in dendritic orientation between control and *βγ*-*cat*^*ChATMN*Δ^ mice (Figure 6b). To assess respiratory circuit function, we performed phrenic nerve recordings in isolated brainstem-spinal cord preparations in control and *βγ*-*cat*^*ChATMN*Δ^ mice at P4, and found that *βγ*-*cat*^*ChATMN*Δ^ mice exhibit similar phrenic MN bursting frequency, burst duration and overall activity as control mice (Figure 6c-d). Our data suggests that cadherins engage catenin signaling during a short temporal window early in development to shape respiratory motor output, but appear to be dispensable once early phrenic MN topography and morphology have been established.

**Figure 6.**
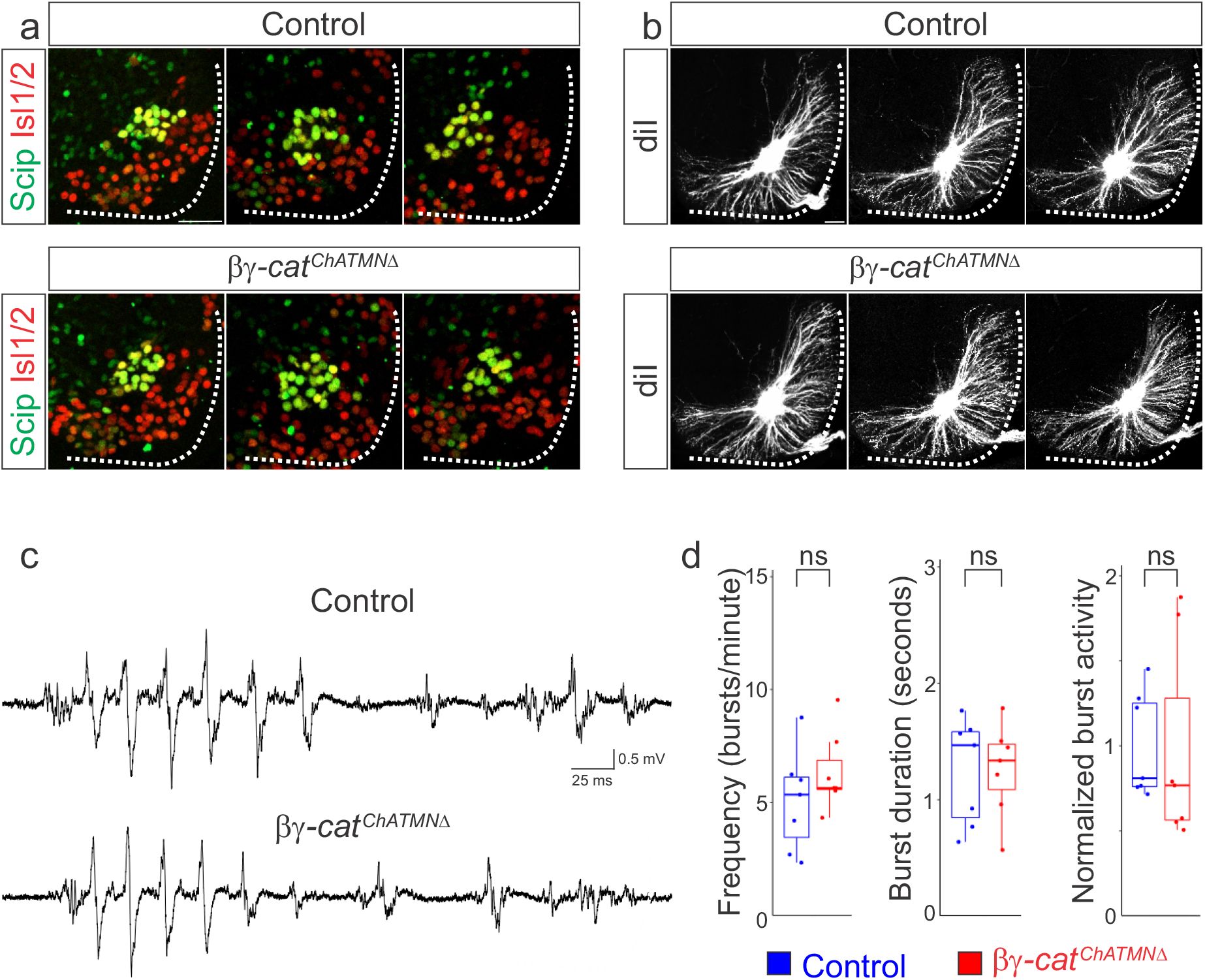
A narrow temporal requirement for catenin signaling in phrenic MN topography and function. **a)** Examples of phrenic MN cell body distribution in control and *βγ*-*cat*^*ChATMN*Δ^ mice at e13.5. *ChAT::Cre*-mediated catenin deletion from postmitotic MNs does not affect their position or clustering. Scale bar= 25μm. **b)** diI injections into the phrenic nerve reveal similar dendritic architecture in control and *βγ*-*cat*^*ChATMN*Δ^ mice at e18.5. Scale bar= 100μm. **c)** Representative suction electrode recordings from the phrenic nerve in P4 control and *βγ*-*cat*^*ChATMN*Δ^ mice show similar levels of phrenic MN activation. **d)** Burst frequency, duration and integrated activity are unchanged in *βγ*-*cat*^*ChATMN*Δ^ mice (n=7 control, n=7 *βγ*-*cat*^*ChATMN*Δ^ mice).

## Discussion

Phrenic MNs are a critical neuronal population that is essential for breathing, yet the molecular mechanisms that control their development and maintenance have remained elusive. Here, we show that catenin-mediated cadherin adhesive function is required for phrenic MN organization, axonal and dendritic arborization, and respiratory output during a narrow developmental window. While catenins have a critical role in early phrenic MN development, they appear to be dispensable for maintaining the morphology and function of these MNs. Our findings indicate that distinct molecular pathways are likely to mediate the establishment and maintenance of respiratory motor circuits at different timepoints throughout development and adulthood.

Catenins appear to be critical for phrenic MN organization. We find that catenin inactivation leads to both ventral shifts in PMC position and loss of clustering between cell bodies. While we observe similar shifts in cell body position when we inactivate 4 out of the 6 cadherins expressed in PMC neurons (cadherins N, 6, 9 and 10-*N*^*MNδ*^*6910^/-^* mice), we do not see a loss of clustering in these mice (Vagnozzi, 2022). This could suggest that retaining expression of the remaining PMC-specific cadherins, 11 and 22, is sufficient to maintain the phrenic MN distinct tight clustering organization. Alternatively, our data could indicate that cell non-autonomous cadherin function plays a predominant role in MN clustering, and that eliminating cadherin signaling from all MNs leads to scattering and mixing of MN populations, causing disorganization not seen when solely eliminating PMC-specific cadherins. Despite differentially affecting PMC clustering, we observe similar changes in phrenic MN activity in *N*^*MNδ*^*6910^-/-^* and *βγ*-*cat*^*MN*Δ^ mice, suggesting that MN clustering may not significantly contribute to phrenic MN activity. Alternatively, since the loss of activity we observe in both mouse models is so severe, it may mask a subtler impact of clustering to PMC activity patterning and synchronization. Decoupling PMC clustering from changes in neuronal morphology and cell adhesion loss will help distinguish the contribution of each of these properties to respiratory motor output.

Our data show that cadherin signaling is required both for the elaboration of PMC axons and dendrites, however the impact of catenin inactivation on dendrites appears to be more severe. Phrenic MNs are able to elaborate axons in *βγ*-*cat*^*MN*Δ^ mice and axonal topography and orientation are mostly preserved, with some minor loss of terminal arborization. Dendrites however appear to be severely stunted, project haphazardly and their topography is lost. This indicates that cadherins have a much more predominant role in dendritic rather than axonal elaboration. While many signaling pathways contribute to phrenic axon growth and diaphragm innervation, including HGF/MET (Sefton et al., 2022), Slit/Robo (Charoy et al., 2017) and Col25a1 (Tanaka et al., 2014), to our knowledge, cadherins are the first cell adhesion molecules to be implicated in phrenic MN dendritic development.

Loss of catenin-mediated cadherin adhesive function results in a dramatic reduction of phrenic MN activity that leads to perinatal lethality. This could be due to the loss of descending inputs from brainstem respiratory centers that provide excitatory drive to initiate diaphragm contraction during inhalation. The dramatic change in PMC dendritic coordinates is likely to contribute significantly to the loss of presynaptic inputs and respiratory activity. Dendrites represent the largest surface area of neurons, and thus receive the majority of synaptic input. In sensory-motor circuits, proprioceptive inputs are primarily located on the dendrites of motor neurons, and different motor pools exhibit distinct, stereotyped patterns of dendritic arborization that contribute to sensory-motor specific connectivity (Balaskas et al., 2019; Vrieseling and Arber, 2006). This mode of cadherin action in respiratory circuits would be consistent with cadherin-dependent targeting mechanisms in the retina, where combinatorial codes of cadherin expression serve to direct axons and dendrites of synaptically connected neurons to their correct laminar targets (Duan et al., 2014; Duan et al., 2018; Osterhout et al., 2011).

In addition to establishing phrenic MN dendritic morphology, cadherins could directly contribute to phrenic connectivity through establishing a molecular recognition program between phrenic MN dendrites and pre-motor axons. Due to their restricted and selective expression in neural populations, cadherins are thought to function in circuit assembly by dictating synaptic specificity. Cadherin expression often reflects the functional connections formed in a circuit, suggesting they may represent a molecular code dictating the formation of selective synaptic connections (Suzuki et al., 1997). Cadherins are expressed on dendrites, axons, and growth cones of developing neurons (Basu et al., 2015) and have been visualized at synapses in both pre and postsynaptic compartments (Bartelt-Kirbach et al., 2010; Benson and Tanaka, 1998; Bozdagi et al., 2000; Fannon and Colman, 1996; Manabe et al., 2000; Suzuki et al., 2007; Uchida et al., 1996; Yamagata et al., 1995). Therefore, cadherins could function to establish respiratory neuron connectivity independently of their role in dictating axonal and dendritic targeting, at the level of the synapse, as it has been described in the hippocampus (Basu et al., 2017; Williams et al., 2011). Future experiments will determine the primary mode of cadherin action in respiratory circuit formation.

While early cadherin inactivation in MN progenitors results in dramatic changes in phrenic MN morphology and activity and leads to perinatal death, cadherin inactivation in postmitotic MNs does not impact respiratory output. This result provides initial evidence supporting a predominant role for cadherins in shaping dendritic orientation, as they appear to be dispensable once phrenic MN morphology has been established. Our findings indicate a model in which cadherins may function to direct the dendrites and axons of pre and postsynaptic neurons to the correct location, while additional cell adhesion molecules dictate synaptic connectivity. In support of this hypothesis, we have identified a number of cell adhesion molecules that are specifically expressed in phrenic MNs but are not required for their morphology. Our results also indicate that distinct mechanisms may function to maintain respiratory circuit integrity after initial formation. Understanding how these critical circuits are maintained in adulthood is essential, as loss of respiratory function underlies lethality in many neurodegenerative diseases such as Amyotrophic Lateral Sclerosis (ALS).

## Acknowledgements

We thank Heather Broihier, Evan Deneris, Ashleigh Schaffer, Helen Miranda, and members of the Philippidou lab for helpful discussions. This work was funded by NIH R01NS114510 to PP, F30HD096788 to ANV, T32GM007250 to ANV/CWRU MSTP, and F31NS124240 to MTM. PP is the Weidenthal Family Designated Professor in Career Development.

